# Longstanding auditory sensory and semantic differences in preterm born children

**DOI:** 10.1101/2023.07.12.548657

**Authors:** Chrysa Retsa, Hélène Turpin, Eveline Geiser, François Ansermet, Carole Müller-Nix, Micah M. Murray

## Abstract

More than 10% of births are preterm, and the long-term consequences on sensory and semantic processing of non-linguistic information remain poorly understood. 17 very preterm-born children (born at <33 weeks gestational age) and 15 full-term controls were tested at 10 years old with an auditory object recognition task, while 64-channel auditory evoked potentials (AEPs) were recorded. Sounds consisted of living (animal and human vocalizations) and manmade objects (e.g. household objects, instruments, and tools). Despite similar recognition behavior, AEPs strikingly differed between full-term and preterm children. Starting at 50ms post-stimulus onset, AEPs from preterm children differed topographically from their full-term counterparts. Over the 108-224ms post-stimulus period, full-term children showed stronger AEPs in response to living objects, whereas preterm born children showed the reverse pattern; i.e. stronger AEPs in response to manmade objects. Differential brain activity between semantic categories could reliably classify children according to their preterm status. Moreover, this opposing pattern of differential responses to semantic categories of sounds was also observed in source estimations within a network of occipital, temporal and frontal regions. This study highlights how early life experience in terms of preterm birth shapes sensory and object processing later on in life.

**Highlights:**

- How very preterm birth affects nonlinguistic auditory processes in school-age is unknown
- We measured auditory evoked potentials to environmental sounds
- Sensory processing differences manifested from 50ms post-stimulus onwards
- Semantic processing differences manifested at 108-224ms post-stimulus
- Classification of preterm status was possible from semantic processing differences

## Introduction

Every year and worldwide, more than 10% of babies are born preterm (PT; i.e. before 37 weeks gestation) (WHO, 2018). Concomitantly, the increased survival rate of preterm born infants also results in more prevalent manifestations of associated sequelae that can be present in infancy or manifest later during childhood or adolescence (Maitre et al., 2020; Maitre et al., 2013). Multiple studies have shown that PT children present more delays in sensory and cognitive development than their full-term (FT) peers (Aarnoudse-Moens, Weisglas-Kuperus, van Goudoever, & Oosterlaan, 2009; Barre, Morgan, Doyle, & Anderson, 2011; Bhutta, Cleves, Casey, Cradock, & Anand, 2002; Dimitrova et al., 2018; Johnson & Marlow, 2011; Turpin et al., 2019). Especially in the early days following birth, the sensory development of the PT born infant can be affected by pathological processes (e.g. due to hypoxia-ischemia and inflammation) and the environment of the NICU that is full of noise, other stressful stimuli and atypical sensory experiences (Maitre et al., 2014; Vandormael et al., 2019). Recent evidence suggests that the resulting sensory processing deficits can contribute to subsequent cognitive difficulties such as fine motor delays, global cognitive impairment, visual processing issues and atypical language development (Spittle et al., 2016; Hövel et al., 2015; Vandormael et al., 2019; Ters et al., 2018; Chorna et al,2014). In turn, this cascade might lie at the core of developmental challenges, school readiness and poorer outcomes later on (Ayres & Robbins, 2005; Maitre et al. 2020; Maitre et al., 2017). Therefore, it is important to characterise any links between low-level sensory and high-level cognitive processing patterns in PT children, as these will likely help to devise appropriate interventions to rehabilitate or reinforce these processes early on.

At birth, the sensory systems of PT infants are immature and thus challenged in appropriately managing the information in the extra-uterine environment (Blackburn, 1998). Moreover, the NICU environment is itself atypical and noisy across multiple sensory modalities, which is considered to contribute to sleeping problems and even alterations of cardiac rhythm in newborns (Wachman & Lahav, 2011). Although contrasting evidence exists (Nevalainen et al., 2008), impaired sensory processing in infancy has been increasingly documented in preterm born infants (Broring et al., 2017; Cabral et al., 2016; Mitchell et al., 2015; Crozier et al., 2015). PT children have a disadvantage when processing sensory information at least in early childhood. Standardized tests suggest atypical developmental patterns across sensory modalities, some of which persist into later childhood (Bucci, Wiener-Vacher, Trousson, Baud, & Biran, 2015; Jackson, Ong, McIndoe, & Ripley, 2003; Jongmans et al., 1996; Wickremasinghe et al., 2013). Recent studies using electroencephalography have shown evidence of atypical processing of tactile (Maitre et al., 2017), auditory (including voice processing; Adam-Darque et al., 2020), and multisensory stimuli (Maitre et al., 2020) in PT children. PT infants have increased risk for hearing loss than FT children and they frequently exhibit auditory processing deficits in early childhood (Gallo et al., 2011). These deficits can be attributed to a combination of the extenuated immaturity of the brain at birth of preterm infants as well as their acoustically atypical environments. This may make them less able to fully benefit from postnatal experience during the first 4 months of life, such as the presence of speech sounds in the environment to which they may have diminished/absent access depending on their NICU experience (Key et al. (2012).

In the auditory domain, hearing deficits have been reported in PT children at 4-7 years of age (Gallo, Dias, Pereira, Azevedo, & Sousa, 2011) and central auditory processing deficits are observed in 7-13 year-old PT children (Amin et al., 2015). Another study of PT infants (6-12 months of age (corrected age)) showed impaired perceptual narrowing (i.e. preferential processing of sounds of their native language) (Jansson-Verkasalo et al., 2010). Auditory processing differences between PT and FT infants and young children have also been documented using auditory evoked potentials (AEPs). These AEP differences manifest not only in terms of amplitude and latency of responses, but also in terms of AEP topography (Mahmoudzadeh et al., 2018; Chorna et al., 2018; Key et al., 2012; Bisiacchi et al., 2009; Fellman et al., 2004; Maitre et al., 2014). AEPs are highly sensitive to differences in sensory processing capacity and can even assess small effects of sensory interventions during infancy (Chorna, Hamm, Shrivastava, & Maitre, 2018). Most previous AEP studies investigated responses to speech stimuli (Chorna et al., 2018; Jansson-Verkasalo et al., 2010; Key, Lambert, Aschner, & Maitre, 2012; Paquette et al., 2015) and/or focused on paradigms eliciting a mismatch negativity (MMN) response. Some evidence would suggest that while speech processing is particularly impaired in PT infants, processing of non-speech sounds (i.e. stimuli based on the 2^nd^ and 3^rd^ formants of the speech stimuli) remains unimpaired (Paquette et al., 2015). Smaller auditory MMN amplitude was observed in 4-year-old children that had been born preterm and with very low birthweight (Jansson-et al., 2003). A similar effect was observed in an auditory distraction paradigm in children at age of 5 years (Mikkola et al., 2010). However, paradigms used in different studies investigating auditory processing in PT children and adolescents have produced inconsistent results. For example, Gomot et al., 2007 showed a normal MMN in 9-year old PT children, albeit reduced amplitude of the later N250 component. These authors hypothesized that this N250 difference may affect development of high-order processes such as language acquisition. Another MMN study on 10-year-old PT children showed shorter N100 latencies and larger P2 amplitudes, suggesting a more stimulus driven response mode (Lindgren et al., 2000). In contrast, Lavoie et al., 1997, showed normal N100 and P2 responses in 5-year-old PT children, but decreased amplitudes of P3a to rare tones. More recently, P300 differences between PT and FT 8-10 year-old children were observed in terms of latency, amplitude and morphology (Durante et al., 2018). More generally, differences in language skills between FT and PT born children have been observed that persist in adolescence (16 years old) (Mullen et al., 2011), albeit diminishing with age (Saavalainen et al., 2006).

The above evidence indicates that auditory processing in PT infants and children differs from that of their FT peers and that such deficits might extend to higher-order auditory processes. Until now, the overwhelming majority of research has not considered to what extent auditory semantic representations of non-speech stimuli – i.e. sounds of environmental objects – are comparable between PT and FT children. Discrimination and categorization of auditory objects is fundamental in everyday life as it englobes everyday interactions that we have with the world (Lewis & Lewis, 2017). Based on evidence in the visual domain, showing differences in processing of animal and tool stimuli between PT and FT adolescents between 13-15 years (Klaver et al., 2015), there is reason to speculate that semantic representations in the auditory modality are also altered by preterm birth.

The aim of the present study is to clarify this and investigate the likely presence of low-level auditory impairments and higher-level cognitive differences in auditory object processing. To this end, 10 year-old children born preterm performed an auditory oddball task involving the discrimination of sounds of living and sounds of manmade objects. This is a well-established task (e.g. Murray et al., 2006) that allows us to explore the interaction between the lower-level sensory and higher-level semantic processes that result in auditory object recognition. AEPs were concurrently recorded in order to compare the brain responses to living and manmade sounds in terms of response strength and topography. Previous studies in healthy adults employing this living/manmade oddball task have shown that differential processing of living versus manmade sounds starts approximately 70ms post-stimulus in terms of response strengths and 150ms in terms of topography (Murray et al., 2006). Furthermore, in terms of topography, two stages in the processing of sounds of living and manmade objects have been demonstrated: an early stage, behavior independent, around 100ms post-stimulus and a later stage linked to behavior and decision making processes, around 270ms post-stimulus (De Lucia et al., 2012). This is the first study to our knowledge investigating auditory object processing in both PT and FT children.

## Methods

### Participants

Thirty-two children participated in the study. The children belonged to two groups: a very preterm group and a full-term group. The VPT group was composed of 17 children (9 female, mean age ±SD = 10.42±0.70 years). VPT children in this sample were recruited from a longitudinal clinical cohort study that investigated how neonatal and parental stress affects children’s development. All VPT children were born at <33 weeks gestational age (GA) (mean GA ±SD = 29.68 weeks±1.92) between 2005 and 2007, and were hospitalized in the NICU of the Lausanne University Hospital. The mother was informed about the study after the child’s birth. In the beginning of the study, infants with malformation or chromosomal abnormalities or parents with psychiatric illness, drug abuse or without fluency in French were excluded. The FT group was composed of 15 children (7 female, mean age ±SD = 10.28±0.98 years). Inclusion criteria were being born at the 37^th^ week or later GA and parental absence of psychiatric illness. FT children were recruited using an advertisement posted in the Lausanne University Hospital and a sports club. Children belonging to the FT group participated only in the current study. No differences between groups were observed in age, gender proportions, or socio-economic status (SES) score (Table 1). SES was assessed based on an adapted version of the Hollingshead Four Factor Index of Socioeconomic status (Pierrehumbert et al., 1996). The total score combined parents’ education level and work position. Higher score means higher SES. For all participants, parental informed written consent was obtained before testing. The procedure was evaluated and approved by the Vaudois Cantonal Ethics Committee (Ethics approval number:256/14).

**Table 1.** Demographic data from the preterm (PT) and full-term (FT) groups as well as their statistical comparison.

|  | PT group | FT group | Statistical test |
| --- | --- | --- | --- |
| N girls | 17 (9) | 15 (7) | $\chi^2 = 0.125, p > 0.1$ |
| Age (years) | 10.42 (0.70) | 10.28 (0.98) | $t_{(30)} = 0.479, p > 0.1$ |
| SES | 2.82 (0.50) | 3.00 (0.65) | $t_{(30)} = -0.871, p > 0.1$ |

### Apparatus and stimuli

The participants were seated at the center of a sound-attenuated booth (WhisperRoom model 102126E), and acoustic stimuli were delivered over inset earphones (Etymotic model ER-4P; www.etymotic.com). Stimulus intensity was approximately 75dB SPL at the ear (measured via a CESVA audiometer). Auditory stimuli were complex, meaningful sounds (16 bit stereo; 22,500Hz digitalization) taken from Murray et al. (2006). The sound set was composed of sounds of living objects and sounds of manmade objects (see Table 2). In total, there were 120 different sound files: 60 sound files of living objects (20 different living objects presented in three exemplars) and 60 sound files of manmade objects (20 different manmade objects presented in three exemplars). Each sound was 500ms in duration including an envelope of 50ms decay time that was applied to the end of the sound file. The E-prime software controlled stimulus delivery and recorded the participants’ behavioural performance (www.pstnet.com/eprime).

**Table 2.** Sound objects used in this study (see fuller details including psychoacoustics in Murray et al., (2006)). Note that sounds of humans were all non-speech vocalizations. Note also that sounds of musical instruments entailed a few notes, but not any melody or rhythms. For all sounds there were three acoustically distinct versions used.

| Sounds of living objects |  | Sounds of manmade objects |  |
| --- | --- | --- | --- |
| Baby crying | Frog | Accordion | Guitar |
| Bird | Gargling | Bicycle bell | Harmonica |
| Cat | Laughter | Car horn | Harp |
| Coughing | Owl | Cash register | Organ |
| Chicken | Pig | Church bell | Piano |
| Clearing throat | Rooster | Cuckoo clock | Police siren |
| Cow | Scream | Doorbell | Saxophone |
| Crow | Sheep | Door closing | Telephone ringing |
| Dog | Sneezing | Flute | Trumpet |
| Donkey | Whistling | Glass shattering | Violin |

### Procedure

Participants performed an auditory ‘oddball’ detection task with living vs. manmade sounds. Every participant completed four blocks. The ‘target’ category of stimuli to which participants responded occurred 10% of the time for a given block. The remaining 90% of stimuli (‘distracters’) were comprised of the other sound category. Therefore, two out of the four blocks had manmade sounds as targets, and two blocks had living sounds as targets. Every block contained 150 trials with a variable inter-stimuli interval ranging between 1500 and 2000ms. As all sounds had duration of 500ms, each block lasted approximately 5 minutes. A short instruction in the screen, accompanied by a more detailed verbal explanation of the task by the experimenter, was presented before the start of each block. In addition, examples of living and manmade sounds were presented to children to verify their understanding of the task at the beginning of the experiment. The children’s task was to press as quickly and accurately as possible a button on the response box when the target stimuli appeared. The same response button was used in all blocks. Children were encouraged to take short breaks between blocks in order to minimize fatigue.

### EEG recording and pre-processing

Continuous EEG was acquired at 1024Hz through a 64-channel Biosemi ActiveTwo AD-box (www.biosemi.com), referenced to the common mode sense (CMS, active electrode) and grounded to the driven right leg (DRL; passive electrode), which functions as a feedback loop driving the average potential across the electrode montage to the amplifier zero. Prior to epoching, the EEG was filtered (low-pass 40Hz; high-pass 0.1Hz; removed DC; 50Hz notch; using a second-order Butterworth filter with -12dB/octave roll-off that was computed linearly in both forward and backward directions to eliminate phase shifts). Peri-stimulus epochs from distracter trials, spanning 100ms pre-stimulus to 500ms post-stimulus onset, were averaged from each subject for each condition to compute AEPs. EEG responses to target stimuli were not analysed as they were too few for sufficient signal quality. Epochs were rejected based on automated artefact rejection criterion of ±80μV as well visual inspection for eye blinks and movement or other sources of transient noise. The average (±SD) number of accepted EEG epochs for the living sounds condition, for FT children was 159±27 and for VPT children was 158±25. The average number of accepted EEG epochs for the manmade sounds condition, for FT children was 164±27 and for VPT children was 156±21. No differences in these values were observed. Neither significant main effect of Group or Category nor the interaction between the two factors was observed (F_(1,30)_=0.26, p=0.69; F_(1,30)_=.13, p=0.78; and F_(1,30)_=0.99, p=0.48, respectively). Bad channels were identified before averaging and excluded from the artefact rejection. These data at artefact electrodes from each participant were interpolated using 3-D splines prior to group averaging (Perrin, Pernier, Bertnard, Giard, & Echallier, 1987). The average number of interpolated channels was 4.8±1.3. In addition, data were baseline corrected using the 100ms pre-stimulus period and recalculated against the average reference.

### AEP analyses

Differences in the processing of AEPs to living and manmade sounds between FT and VPT children were assessed using a multi-step analysis procedure, referred to as electrical neuroimaging, which involves both local and global measures of the electric field on the scalp. These methods have been described in detail previously (Koenig, Stein, Grieder, & Kottlow, 2014; Michel & Murray, 2012; Michel et al., 2004; Murray, Brunet, & Michel, 2008; Tzovara et al., 2012).

First, we analysed the AEP voltage waveform data from each scalp electrode as a function of time using a two-way mixed-model repeated measures analysis of variance (rmANOVA) with the between-subject factor of children group (2 levels: FT vs. PT) and the within-subject factor of type of semantic Category (2 levels: Living vs. Manmade). For this analysis we used an average reference as well as a temporal criterion for the detection of statistically significant effects (>10ms continuously at 1024Hz sampling rate) in order to correct for temporal auto-correlation at individual electrodes (Guthrie & Buchwald, 1991). Similarly, a spatial criterion (effects were considered statistically significant only if they entailed >10% of the electrodes of the 64-channel montage at a given latency) was applied in order to address spatial correlation.

Second, a topographic cluster analysis based on a hierarchical clustering algorithm was performed on the post-stimulus group-average AEPs across experimental conditions (Murray et al., 2008). This clustering (“segmentation”) identifies stable electric field topographies (“template maps”). The clustering is insensitive to pure amplitude modulations across conditions as the data are first normalised by their instantaneous GFP. The optimal number of template maps that explained the whole group-averaged data set was determined using a modified Krzanowski-Lai criterion (Murray et al., 2008). The clustering makes no assumption regarding the orthogonality of the derived template maps (De Lucia et al., 2010; Koenig et al., 2014; Pourtois, et al., 2008). The pattern of maps that was identified in the group-average AEPs was then submitted to a fitting procedure wherein each time point of each individual participant’s ERP is labelled according to the template map with which it best correlated spatially (Murray et al., 2008). This yielded a measure of relative map presence over a fixed time window (in milliseconds) that was then submitted to a mixed-model rmANOVA with factors of Group, Map, and Category. This fitting procedure revealed whether a given experimental condition for a given group was more often described by one map versus another, and therefore whether different intracranial generator configurations better accounted for particular experimental conditions or groups.

Subsequently, changes in the strength of the electric field at the scalp as a function of semantic category and group of children were assessed using global field power (GFP) for each participant and condition (Lehmann, 1987). GFP is equivalent to the standard deviation of the voltage potential values across the electrode montage (Lehmann & Skrandies, 1980). A stronger GFP value is indicative of greater and/or more synchronised brain activity, though the root cause (increased neural firing rate, increased numbers of active neurons, etc.) cannot be unequivocally asserted based on this measure alone. However, a modulation of GFP in the absence of reliable evidence for topographic modulations can most parsimoniously be interpreted as a modulation in the strength of responses originating from statistically indistinguishable sources or set of sources. GFP data were analysed using area measures. These measures were calculated (vs. the 0µV baseline) using time periods of stable scalp topography that were defined in the previous analysis steps and were statistically tested using rmANOVAs to investigate the effects of Group and Category. Next, we assessed whether differences between semantic categories in GFP area values over an identified time period could reliably classify participants according to their group assignment (PT vs. FT). To do this we conducted a receiver operating characteristic (ROC) and area under the curve (AUC) analysis. ROC analyses were performed in SPSS.

Finally, we estimated the underlying intracranial sources of the AEPs in response to the two different types of sound in the two groups of children using a distributed linear inverse solution (minimum norm) combined with the LAURA (local autoregressive average) regularisation approach (Grace de Peralta Menendez et al., 2001, 2004; see also Michel et al., 2004 for a comparison of inverse solution methods). The solution space was calculated on a realistic head model that included 3000 nodes, selected from a grid equally distributed within the grey matter of the Montreal Neurological Institute’s average brain (available from https://sites.google.com/site/cartoolcommunity/downloads). The head model and lead field matrix were generated with the Spherical Model with Anatomical Constraints (SMAC; Spinelli et al., 2000 as implemented in Cartool version 4.10 (Brunet et al., 2011)), using a 4-shell model (skull, scalp, cerebral spinal fluid, and brain) as well as with an upper skull thickness (5.1mm) and mean conductivity (0.027S/m) values based on the mean age of our sample. As an output, LAURA provides current density measures; their scalar values were evaluated at each node. Statistical analysis of source estimations was performed by first averaging the AEPs across time for each participant and condition over the first and second time windows emerging from the analyses described above. Specifically, during the first time window there was a main effect of Group, and therefore the source modelling was performed after first collapsing the AEP data across factor of Category. For the second time window there was a reliable Category x Group interaction in the ERP analysis and thus a rmANOVA was performed on the source modelled data. The statistical significance criterion at an individual solution point was set at p<.0.05. Only clusters with at least 10 contiguous significant nodes were considered reliable in an effort to correct for multiple comparisons and was based on randomization thresholds (see also De Lucia et al., 2012; Knebel and Murray, 2012; Retsa et al., 2018; Retsa et al., 2020 for similar implementations).

## Results

### Behavioural results

Children in both FT and VPT groups performed the task accurately with no significant main effects of Group (F_(1,30)_=0.49; p=0.492; η^2^=0.144) or Category (F_(1,30)_=3.69; p=0.064; η^2^=0.461), nor an interaction between factors (F_(1,30)_=0.844; p=0.366; η^2^ =0.144; see Table 3). The results of the mixed rmANOVA on the reaction times showed no main effect of Group (F_(1,30)_=0.46; p=0.500; η^2^ =0.101) nor an interaction between Group and Category (F_(1,30)_=3.81; p=0.060; η^2^ =0.472. However, a main effect of Category was observed (F_(1,30)_=46.38; p<0.002; η^2^ =1.0). Overall RTs to manmade objects were faster than RTs to living objects (917ms vs. 1032ms). This difference in RTs was observed for both FT (t_(14)_ = 5.02, p<.001) and PT children (t_(16)_= 4.5, p<.001) and is consistent with prior observations in healthy adults (Bergerbest et al., 2004; Murray et al., 2006; Saygın et al., 2003).

**Table 3.** Group-averaged accuracy and reaction time for each group and each semantic category. Asterisks indicate significant differences between groups.

|  | PT group | FT group |
| --- | --- | --- |
| *Living* |  |  |
| Accuracy (sd) [%] | 91.55 (5.6) | 91.3 (5.1) |
| Reaction time (sd)<br>[ms] | 1029.14 (96.71)* | 1034.96(131.01) |
| *Man Made* |  |  |
| Accuracy (sd) [%] | 93.05 (9.8) | 95.64(2.79) |
| Reaction time (sd)<br>[ms] | 946.85(122.51)* | 886.54(138.56) |

### AEP results

Group-averaged AEPs from FT and VPT children in response to sounds of living and manmade objects from an exemplar midline electrode (FCz) are displayed in Figure 1. The results of the two-way mixed-model ANOVA across the full electrode montage as a function of time showed significant main effects of both children’s Group and sound Category, starting as early as 50ms post-stimulus onset (see Figure 1). The main effect of Group followed from the generally larger magnitude of responses in the FT group than VPT group. There was a general difference between responses to Category starting at 50ms post-stimulus. In addition, there was a significant interaction between Group and Category ∼140-250ms post-stimulus.

**Figure 1.**
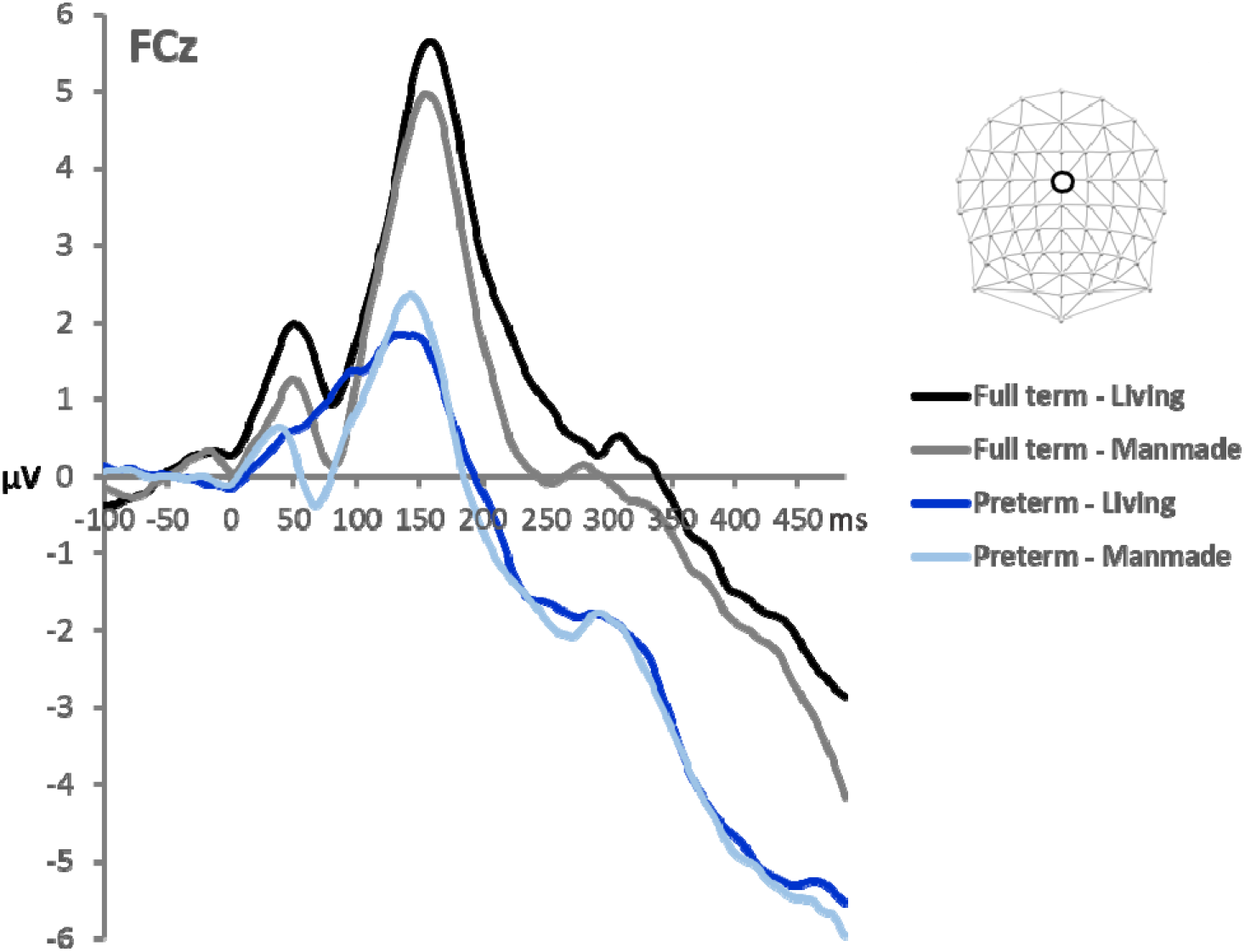
**A.** Group-averaged AEPs at an exemplar fronto-central midline electrode, shown separately for each group and sound-type.

A topographic clustering analysis was then conducted over the full 500ms post-stimulus time period in order to identify time intervals of stable electric field distributions at the scalp and to determine whether the above response differences between conditions followed from single or multiple topographic configuration changes. This analysis provides a set of so-called “template maps”. Eight different template maps (shown in Figure 2) accounted for the collective group-averaged dataset with a global explained variance of 95.5% across the cumulative group-averaged data. During the first interval (36-107ms), two template maps were identified, during a second interval (108-224ms) four maps were identified, and during a third time interval (225-400ms) three template maps were identified. Across the different time intervals shown, the same set of topographies was observed for both living and manmade sounds. That is, the same maps appeared to represent the responses to living sounds and manmade sounds during all the time periods. In contrast, across all the three time intervals (36-107, 108-224, 225-400ms), different maps appeared to predominate the responses of the FT group and the responses of the VPT group. This pattern observed in the group-averaged AEPs was statistically assessed in the single-subject data using the spatial correlation based fitting procedure. This was done separately for each of the three different time periods and their respective template maps. The values of the fitting procedure were then submitted to a mixed rmANOVA (one for each time period) using Group, Category and template Map as factors (see Figure 2 for the number and topography of template maps identified over each time period).

**Figure 2.**
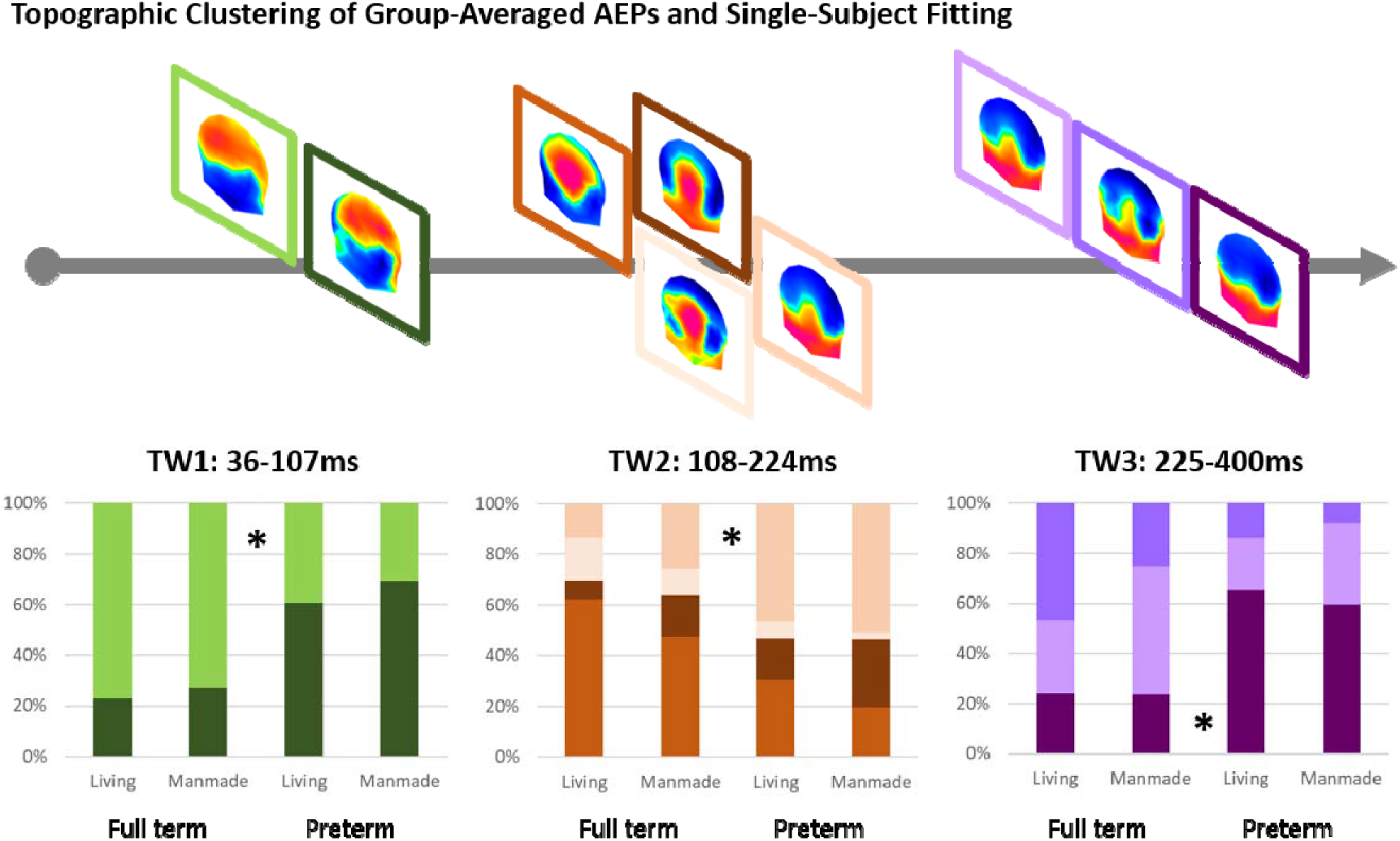
The topographic pattern analysis identified eight stable topographies (template maps) for all the conditions around 500ms post-stimulus period. The time period when each map was observed is indicated. At the group-average level, two different template maps were identified over the 1) 36-107ms post-stimulus period, four different template maps over the 2) 108-224ms period and three maps over the 3) 225-400ms post-stimulus period. The bar graphs on the bottom show the results of the single-subject fitting procedure, indicated the group-averaged duration each template map was ascribed to each of the two groups. All the three mixed ANOVAs (one for each time period) on these duration values revealed an interaction between template map and group. Post-hoc comparisons in the all the three time periods demonstrated the maps that are predominant for each group.

For the first time period (TW1=36-107ms), there was a significant interaction between Group and Map (F_(1,30)_= 8, p< .001, *η_p_^2^*=0 .211). That is, one map predominated the responses of the FT group and the second map predominated the responses of the VPT group (see Figure 2).Also, over the second time window (TW2=108-224ms), there was a significant interaction between Group and Map (F_(3, 90)_ = 5.8, p<.001, *η_p_^2^*=0 .162), indicating that different template maps predominated responses in each of the two groups. As there was no interaction between Category and maps nor a three-way interaction, we averaged the data across responses to living and manmade sounds for each map in order to test which maps predominantly represent the FT and the VPT groups. The results of this analysis showed that one template map predominated the FT group and another map the VPT group (see Figure 2). Over the third time period (TW3=225-400ms), there was again an interaction between Group and Map (F_(2,60)_= 6.64, p<.01, *η_p_^2^*=0 .181). Similarly to the second time window, the data from living and manmade semantic categories were collapsed in order to investigate the contribution of the three template maps on the two groups. Two template maps found to predominate responses from the FT group, whereas the third map was found to predominate responses from the VPT group (see Figure 2).

Prior data from adults indicates that semantic processing of sounds of objects involves a latency difference expressed as the difference in the duration over time when a given map offset (i.e. the last time point when a template map best correlated spatially with AEPs from each semantic category) (Murray et al., 2006). We therefore performed a similar analysis here using the last offset of template maps over the 108-224ms post-stimulus period. We observed a significant three-way interaction between Group, Category and Map (F_(3,87)_ = 9.65, p< .001, *η_p_^2^*=0 .286). Post-hoc comparisons showed that there was an effect of Category only in the FT group. Specifically, the predominant map for FT in this time window was found to end significantly later in response to living sounds (204ms±0.76(SE)) compared to manmade sounds (182ms±3.7(SE)).

Based on the results of the topographic clustering analysis, we also defined the time windows for the analysis of GFP waveforms (Figure 3A). Specifically, we investigated the main effects of Group and Category as well as their interaction on GFP area for each of the three identified time windows. Neither the main effect of Group nor Category was found to be significant in any of the three time windows. The Group × Category interaction was significant only for TW2 (108-224ms): F_(1,30)_= 11.4, p<.01, *η_p_^2^*=0 .275. Post-hoc t-tests indicated that in the FT group children responded significantly stronger to sounds of living objects (2.87μV) compared to sounds of manmade objects (2.4μV): t_(14)_=2.83, p<.05. In contrast, in the VPT group, there was a trend for the children to respond stronger to the sounds of manmade objects (2.85μV) than to the sounds of living objects (2.55μV): t_(16)_=1.992, p=.055. (Figure 3B).

**Figure 3.**
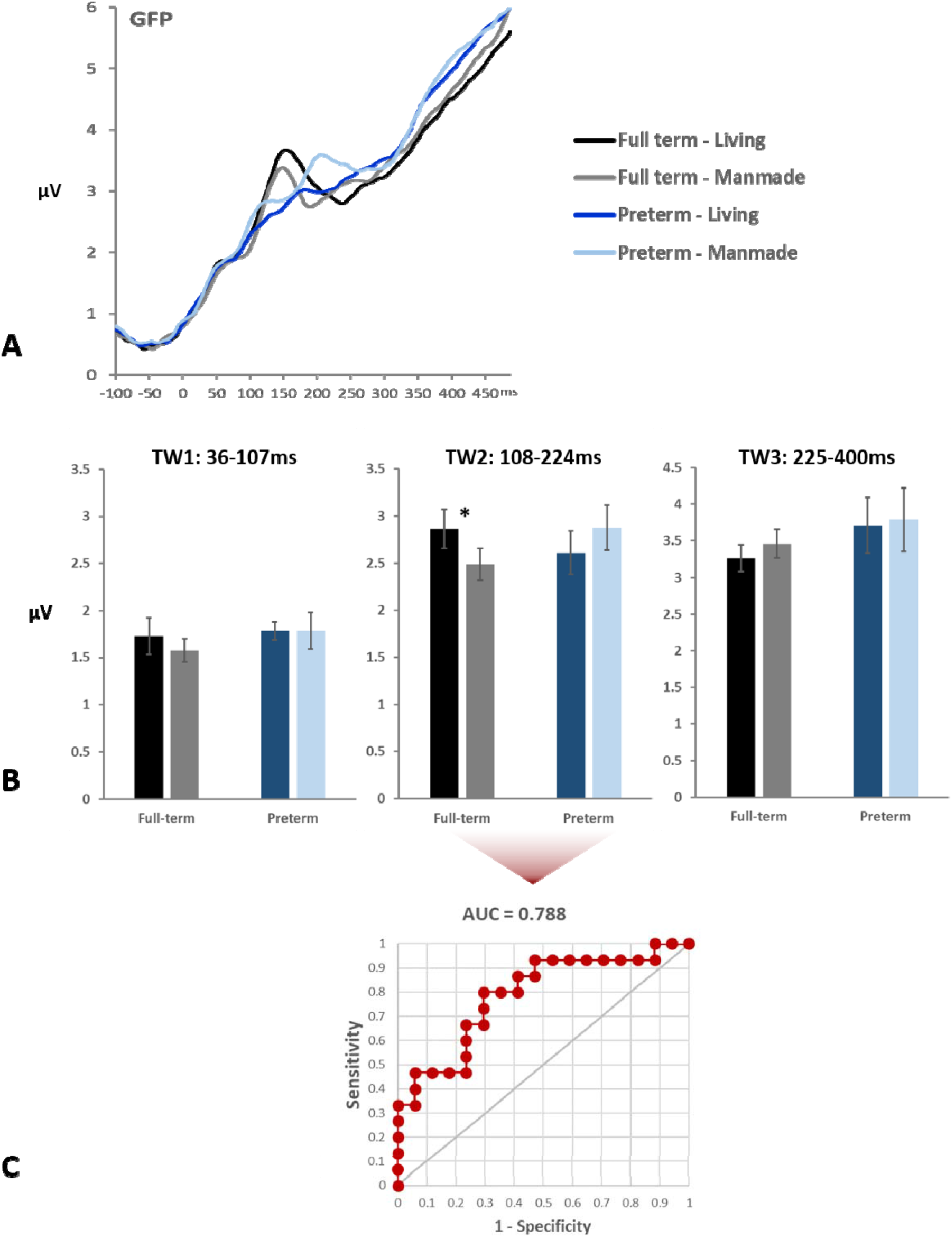
**A.** Global field power (GFP) waveforms from each group and sound-type. **B.** Mean GFP values shown separately for each group and type of sound object in the three identified time periods during the topographic analysis. There is an interaction between group and type of object only in TW2. Asterisks indicate significant differences between types of objects within a group. **C.** ROC curve with AUC score for the GFP differences between living and manmade objects during TW2.

Next, ROC analysis was performed, using the area under the curve (AUC) versus a null hypothesis of chance classification, to determine if VPT and FT children could be reliably classified based on the GFP area measures over TW2, which also exhibited a significant Group x Category interaction (i.e. when we observed differences between the processing of living and manmade objects as a function of prematurity). The ROC analysis showed statistically reliable classification of VPT from FT children, using the differences between the GFP of responses to living and manmade objects, with an AUC of 0.788 (see Figure 3C). In other words, differences in brain responses to semantic categories were sufficient to reliably classify schoolchildren as having been born preterm or full-term.

Finally, LAURA-distributed source estimations were calculated separately over TW1 (36-107ms) and TW2 (108-224ms). For this purpose, AEPs for each participant and experimental condition were averaged over the periods 36-107ms (TW1) and 108-224ms (TW2). For TW1, only overall differences between the two groups of children were investigated. This t-test showed significant differences between FT and PT children in the estimated source activity within the left inferior lingual gyrus (local maximum: -4,-80,-4mm (BA18)), the left superior temporal gyrus (local maximum: -57, -19, 4mm (BA41)), the left dorsolateral prefrontal cortex (local maximum: -50, 27, 19mm (BA46)) and the right anterior prefrontal cortex (local maximum: 11, 57, -11mm (BA10) (Figure 4). Stronger activity in PT children was observed only in the occipital cortex cluster. In the other three clusters, FT children exhibited stronger activity.

**Figure 4.**
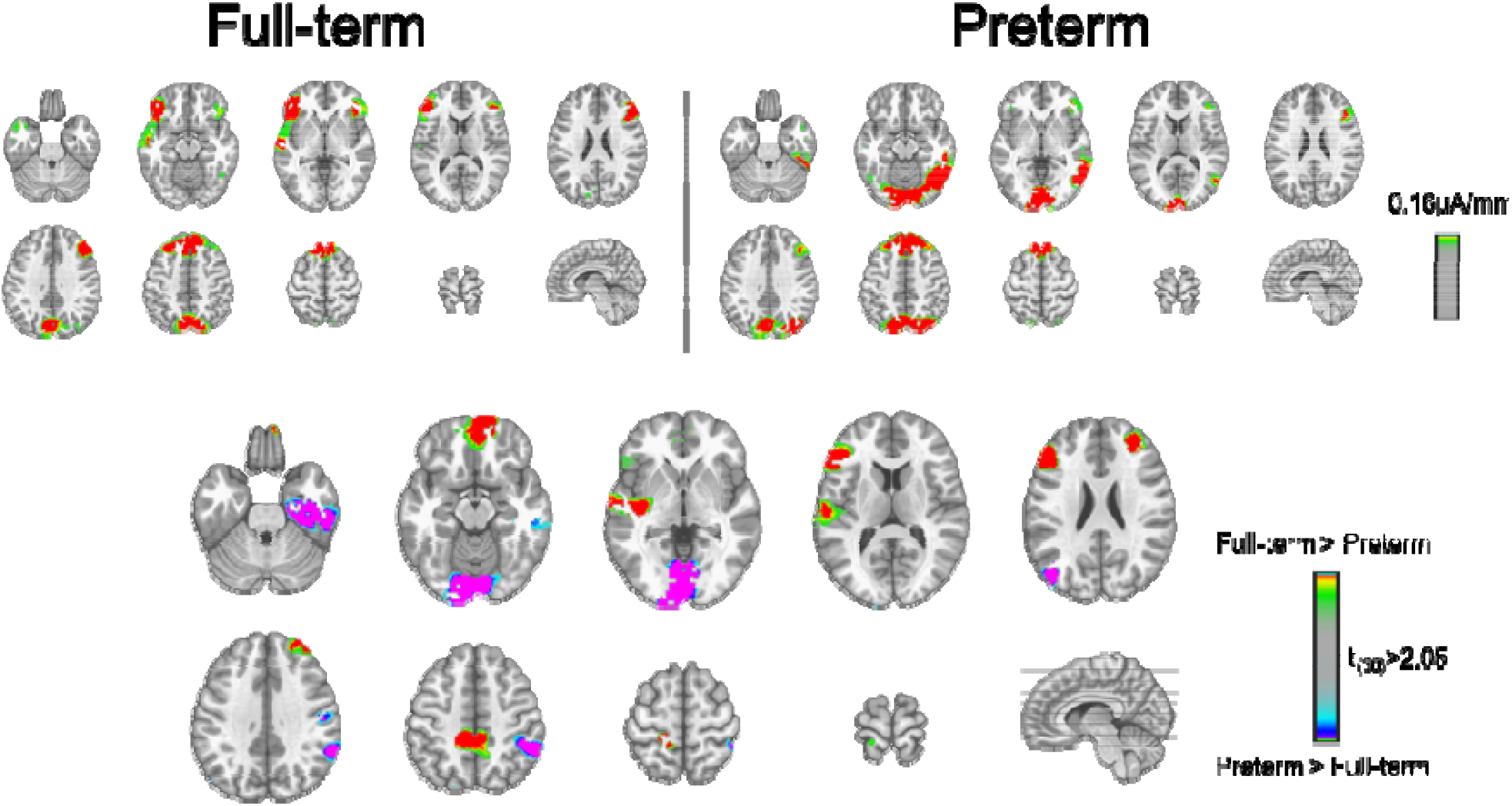
LAURA source estimations over the **TW1 (36-107ms)**. Mean source activations are shown for Full-term and Preterm children independently of semantic group. Significant differences between the two groups of children were observed in four main areas. Stronger activation in the left inferior occipital cortex is observed in PT children, whereas stronger activation in the left primary auditory cortex, left dorsolateral prefrontal cortex and right anterior prefrontal cortex was observed in Full-term children.

For TW2, differences between categories and groups were investigated. An rmANOVA was performed and it revealed a significant interaction between Group and Category on the estimated source activity. Differential source activity was localised to clusters within the right inferior occipital cortex (lingual gyrus, local maximum: 25, -72, -2mm (BA 19)) and the right dorsolateral prefrontal cortex (local maximum: 19, 42, 31mm (BA 9)) (Figure 5). Greater differences in the activity between living and manmade objects in the right occipital cortex were observed in PT children, whereas greater differences in the activity between living and manmade objects were observed in FT children in the right prefrontal cortex.

**Figure 5.**
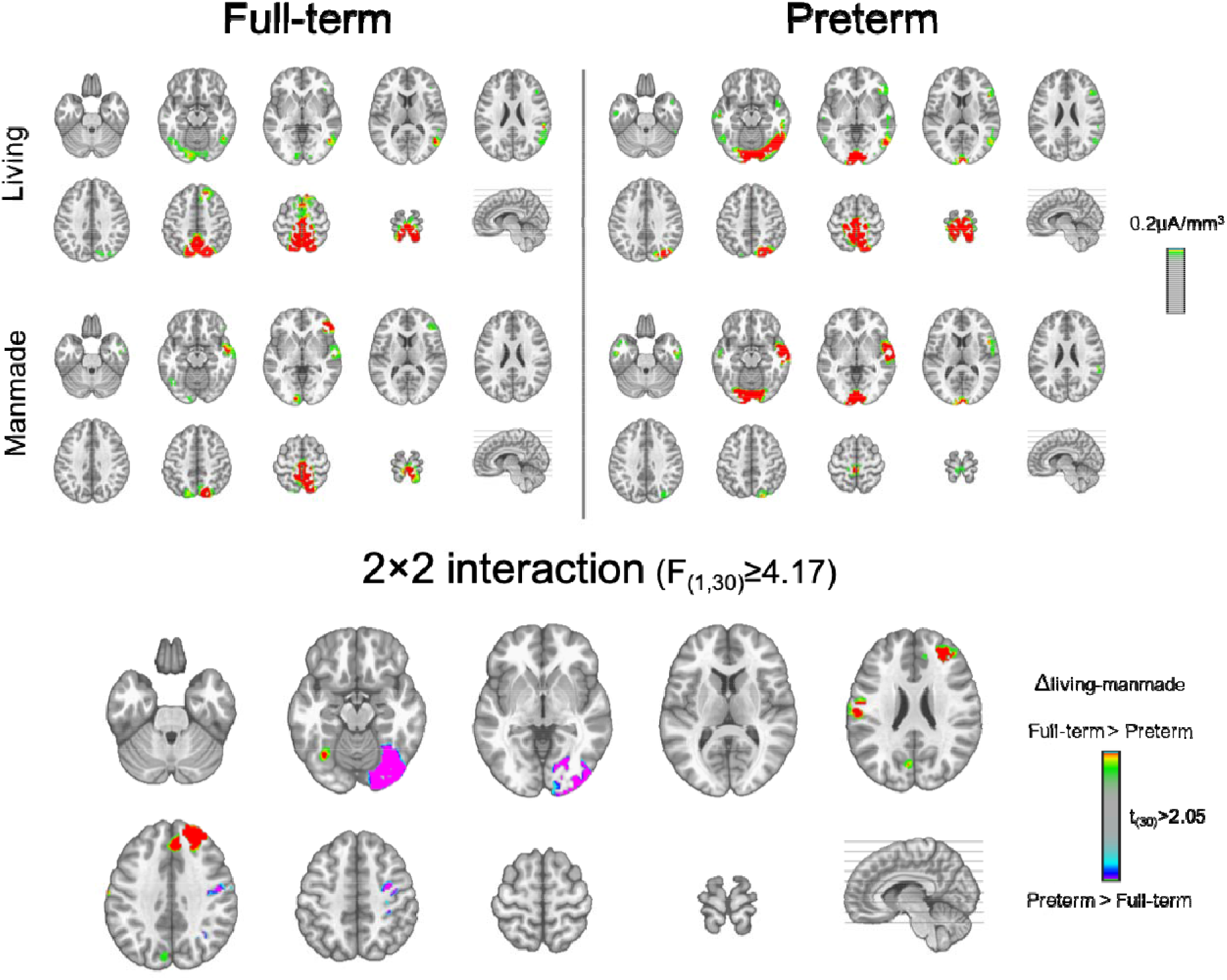
LAURA source estimations over the **TW2 (108-224ms)**. Mean source activations are shown for Living and Manmade objects in Full-term and Preterm children. A significant Group × Condition interaction is observed in two main areas. Greater activation in the right inferior occipital cortex is observed in Preterm children, whereas greater activation in the right superior frontal gyrus is observed in Full-term children.

## Discussion

We investigated the long-term effects of preterm birth on sensory and semantic auditory object processing. An electrical neuroimaging analysis framework applied to the recorded AEPs allowed us to differentiate between effects due to modulations in response strength and effects due to modulations in response topography. We observed general topographic differences in auditory responses between groups as early as 50ms post-stimulus onset, indicative of altered sensory responsiveness and more specifically the engagement of at least partially distinct brain networks by each group in the processing of sounds. Source reconstructions over the 36-107ms post-stimulus period identified differential patterns of activation between FT and VPT children within the primary auditory cortex and prefrontal cortices as well as the occipital cortex. Over the 108-224ms post-stimulus period, we also observed different patterns of semantic discrimination as a function of preterm status. Full-term children exhibited stronger responses to sounds of living than manmade objects, whereas preterm children exhibited the reverse. This index of semantic discrimination accurately classified a child’s preterm status. Finally, source reconstructions over the 108-224ms post-stimulus period identified interactions between Group and Category within brain circuits also typically implicated in visual object discrimination, consistent with the notion of multisensory, if not altogether amodal and multisensory object representations whose functional organization is impacted in a long-term manner by preterm birth. Collectively, our results indicate there to be persistent differences in sensory and semantic processing in healthy preterm-born children.

We provide evidence that preterm birth can entail longstanding and persist effects (at least) until 10 years of age. We also provide some of the first evidence that differences in auditory information processing extend beyond linguistic stimuli to include the semantic analysis of sounds of environmental objects. These sequelae are undoubtedly the consequence, at least indirectly, of both the atypical auditory sensory experiences of the NICU, which are noisy, together with the immaturely developed sensory systems of the PT born infant (Vitale et al., 2021;Carvalhais et al., 2017; Pineda et al., 2017). Another set of contributing factors likely includes the family environment and in particular the reactions of the parents to the preterm birth, including their stress and consequent manner of interacting with their child (e.g. Muller-Nix et al., 2004; Bozkurt et al., 2017; Turpin et al., 2019).

Several ERP studies in infants have reported differential auditory processing in PT versus FT infants (e.g. de Regnier, 2008; Maitre et al., 2013; Kea et al., 2012; Maitre et al., 2014; Bisicacchi et al., 2009; Fellman et al., 2004). Most of these have shown impaired processing of speech stimuli (e.g. Maitre at al., 2013; 2014), including speech-elicited mismatch negativity (MMN) responses that are either absent (e.g. Fellman et al., 2004) or followed by a decreased N200 component (Bisiacchi et al., 2002). In particular, one study demonstrated impaired processing of complex consonant sounds both in terms of AEP amplitude and also topography, indicative of differences in the underlying neural processes for speech sounds between FT and PT infants (Key et al., 2012), starting mainly from 400ms post-stimulus. By contrast, vowel discrimination was found unimpaired (Key et al., 2012). These differences in the processing of consonant sounds between FT and PT infants were interpreted as an additional requirement of the engagement of attentional processes in the case of PT infants whose sensory system is more immature and struggles more with the accurate discrimination of more complex sounds. It should be noted that early auditory responses (before 150ms post-stimulus) were not investigated.

Most of the studies investigating auditory processing in PT children have focused on infancy and early childhood. Very few studies have investigated the long-term effects of preterm birth on sound processing. Two studies indicated that the auditory processing in 9-year-old PT children differs from that of their full-term peers (Gomot et al., 2007; Korpilahti et al., 2016). Gomot et al. observed an unimpaired MMN in the 9-year-old PT children, but a reduced N250 instead. They interpreted this reduction as indicative of dysfunction in auditory processing as a result of delayed brain maturation in PT infants. Korpilahti et al., observed impaired N200 and N400 in the PT group in response to auditory words and pseudowords. Additionally, these differences in N200 in PT children correlated with auditory attention deficits.

Our results showed that despite the absence of behavioural performance differences between FT and VPT children, there are striking neurophysiological differences in auditory processing between the two groups. These sensory processing differences were identified starting as early as 50ms post-stimulus onset, independently of the semantic category of the sound object being heard. VPT children had generally smaller amplitude of AEP responses and exhibited differences in terms of AEP topography. Topographic differences between FT and VPT children were identified throughout the post-stimulus period. That is, different maps – different cortical networks-represented the responses of VPT children compared to the responses of FT children. VPT born children, even 10 years after birth, exhibit fundamental differences in the way that they process sounds.

This is the earliest post-stimulus difference in terms of auditory processing that has been identified so far in PT born schoolchildren. Previous studies have identified later differences starting around 200ms post-stimulus (Korpilahti et al., 2016; Key et al., 2012; Gomot et al., 2007). However, these prior studies used different analysis frameworks (i.e. only voltage waveform analyses in contrast to our electrical neuroimaging analyses) and stimuli and did not focus on early sensory processing. Source modelling of the early AEP differences identified group-wise effects within a network of temporal, occipital and prefrontal regions. Activations within the superior temporal gyrus and prefrontal cortices have been previously shown to be linked to auditory processing, sound discrimination and category discrimination (Husain & Horwitz, 2006). Specifically, activations within superior temporal gyrus and bilateral prefrontal cortices have been observed in a MEG study investigating the processing of natural and manmade sounds (Salvari et al. 2019). Activations within the superior temporal gyrus and prefrontal cortices network were mainly found in FT children compared to VPT children suggesting long-term effects of preterm birth in these regions. In contrast, significantly stronger activations within left inferior occipital cortices were observed in VPT children comparted to FT children. In agreement, a recent fMRI study has indeed shown ventral occipital cortex activation associated with sounds of objects (as well as images of objects), suggesting that this region can be activated in object processing independently of modality Mattioni et al., 2020; Amedi et al., 2005). Interestingly in our study this occipital activation is observed quite early (already at the first time window(36-107ms post-stimulus) and is more evident in VPT than FT children. One possibility, therefore, is that stronger occipital activity in VPT children could reflect a compensatory mechanism where the auditory processing related areas seem to respond more weakly.

Aside from sensory processing differences, we also observed general, group-independent differences in semantic processing. Semantic discrimination of living versus manmade objects was observed starting as early as 50ms post-stimulus, which is comparable to the findings observed in adults (Murray et al., 2006; De Lucia et al., 2010). Additionally, we observed a subsequent Group × Category interaction, indicating differential processing of living and manmade sounds between FT and VPT children. More specifically, FT children exhibited larger amplitudes of GFP in response to living objects, whereas VPT children exhibited larger amplitude of responses to manmade objects in a time window approximately between 100-200ms post-stimulus (Figure 3). It is interesting to note that the differential pattern we observed here in VPT resembles what has been previously reported in adults (Murray et al., 2006). This suggests that in the typical developmental trajectory, responses are initially stronger in response to living objects and at some point, that reverses. Indeed, FT infants show a preference for voices compared to other sounds, whereas the same is not observed in PT infants (Therien et al., 2004). It would be interesting to identify when (and why) this switch happens as well as what is driving this earlier shift in PT children. One speculative possibility is that PT children have stronger responses to manmade objects in part because of their early life exposure to sounds and noises of machines during their stay in NICU (Carvalhais et al., 2017; Therien et al., 2004). A corollary of this speculation would be that such experiences result in augmented representations, perhaps via statistical learning or similar mechanisms, of the kinds of objects to which the preterm infant is exposed; notably sounds of manmade objects. In this regard, it is noteworthy that we could reliably classify a child’s preterm status based on their differential responses to semantic categories (see Figure 3C). It will likely be informative for future research to ascertain to what extent this may also manifest at earlier ages and effectively indicate the exposure of the child and potential efficacy of any remediation through exposure or experience with more typical sensory environments (see e.g. Chorna et al., 2014 for the example of suckling-driven auditory feedback of the infant’s mother’s voice).

There was no evidence for category-wise topographic differences between the processing of living and manmade sounds in either FT or VPT children. Previous data from healthy adults have identified different AEP topographies that characterize responses to sounds of living and manmade objects during the 155-257ms post-stimulus window (Murray et al., 2006). In this regard, and in contrast to the abovementioned GFP results, the topographic patterns in children remain (somewhat) immature. More specifically, whereas distinct brain networks contribute to adults’ semantic processing of environmental sounds, strength modulations within a statistically indistinguishable brain network account for children’s semantic processing. Nonetheless, we did also observe temporal offset modulations in the pattern of AEP topographies only in FT children, which resembles the temporal pattern observed in adults (Murray et al., 2006). Specifically, the predominant AEP topography over the 108-224ms post-stimulus time window in FT children exhibited a protracted offset in response to living versus manmade sounds of objects. No such temporal difference was observed for VPT children.

With regard to the implicated brain circuits in semantic processing of environmental sounds, we observed a common network of brain areas responding to both categories of sounds and both groups of children. This network included regions of the right temporal cortex, the right dorsolateral prefrontal cortex and the right inferior occipital cortex that are active over the 108-224ms post-stimulus window (see Figure 5). Source activity within the right inferior occipital cortex exhibited larger differences between living and manmade objects in VPT children than in FT children. In contrast, within the right dorsolateral prefrontal cortex, we observed significant differences in source strength between living and manmade objects for FT children only. The differences between living and manmade objects within the identified frontal regions (as well as the temporal one) are consistent with previous studies in healthy adults (Murray et al., 2006; Husain & Horwitz, 2006; Lewis et al., 2005; Lewis et al., 2004; Salvari et al., 2019). Of particular note is the involvement of visual cortices in the representation of sounds of auditory objects, particularly in VPT children, which was also observed over both the 108-224ms as well as the earlier 36-107ms post-stimulus time window (see Figures 4 and 5). However, studies have shown that the ventral occipital-temporal cortex (VOTC) is involved in the representation of different categories of auditory objects in a similar way to their visual counterparts (Mattioni et al., 2020;Amedi et al., 2005). In the case of visual cortical involvement in the auditory object recognition, several propositions have been advanced that are not mutually exclusive. On the one hand, it has been proposed that object representations in visual cortices are multisensory in nature (Murray et al., 2004; Murray et al., 2005; Mahon et al., 2009). On the other hand, it has been proposed that auditory object recognition may involve mental imagery in terms of visualizing the referent object (Lewis et al., 2005). Which specific processes or recognition strategies are operating in FT and PT children will require additional research, but may benefit from analytical approaches such as representational similarity analysis (e.g. Tovar et al., 2020).

Employing our paradigm and studying auditory object processing in both younger PT children and teens as well as adults would be critical for understanding the developmental trajectory of auditory object processing and how it is affected by preterm birth. The identification of long-lasting effects of preterm birth in early auditory processing and auditory object processing is a first step in understanding the neurophysiological mechanisms underlying the difficulties that PT children encounter. Here we demonstrated important differences in early auditory processing in general as well as differences in processing of different semantic categories between FT and VPT 10-year-old children. An optimal sensory system is essential in order to process information from the surroundings and to have a favourable cognitive development. The next step is to link the sensory and semantic processing differences in PT children we identified here to measures of cognitive performance and school success. Korpilahti et al. 2016 for example showed differential processing of words and pseudowords only in 9 year old PT children and not FT children; this distinct pattern of processing in PT children was furthermore linked to lower cognitive performance. To the extent that similar associations can be assayed with non-linguistic stimuli, like those of the present study, will not only allow for studying children from a more diverse range of backgrounds but also studying children at earlier neurodevelopmental stages.

The investigation of long-term impacts of preterm birth on developmental outcomes is essential, considering the increased number of preterm birth survivors and the associated risks and difficulties for preterm born infants that often persist to childhood and affect school performance (Maitre et al., 2020; Maitre et al., 2013; Turpin et al., 2019). Therefore, the identification of the potential difficulties combined with the development of strategies in order to tackle them as early as possible is crucial. For example, recent studies have shown that preterm birth negatively affects school readiness as it increases the risk for various cognitive and sensory abilities difficulties (Taylor et al., 2022; Maitre et al., 2020). PT children have lower IQ scores compared to FT children (albeit in the normal range), and this persists even when tested in later childhood (Kim et al., 2021) or early adolescence (Turpin et al., 2019). PT children often exhibit school difficulties and lower academic achievement levels. Preterm born children present more cognitive, emotional and behavioral difficulties (e.g. anxiety, inattention, and ADHD) and are at a higher risk for internalizing and externalizing behaviors, including during early childhood, compared to their full-term born peers (Taylor et al., 2022; Maitre et al., 2020; Aarnoudse-Moens, Weisglas-Kuperus, van Goudoever, & Oosterlaan, 2009; Barre, Morgan, Doyle, & Anderson, 2011). Preterm birth is also considered a high risk factor for subsequent development of neuropsychiatric disorders such as ADHD, inattention, anxiety, autistic spectrum disorders as well as increased risk for psychosis, depression and bipolar disorder (Nosarti et al., 2012; Johnson & Marlow, 2011).

## Author Contributions (CRediT)

Conceptualization and Methodology: CR and MMM; Formal analysis: CR, HT, EG and MMM; Writing of original draft: CR, HT, EG, FA, CMN, and MMM; Supervision: CR, CMN, and MMM).

## Acknowledgements

This work has been financially supported by The Swiss National Science Foundation (grants PZOOP1-148184 to EG, 320030-169206 to MMM)

## References

Aarnoudse-Moens, C. S. H., Weisglas-Kuperus, N., van Goudoever, J. B., & Oosterlaan, J. (2009). Meta-analysis of neurobehavioral outcomes in very preterm and/or very low birth weight children. Pediatrics, 124(2), 717–728.

Amedi, A., von Kriegstein, K., van Atteveldt, N. M., Beauchamp, M. S., & Naumer, M. J. (2005). Functional imaging of human crossmodal identification and object recognition. Experimental brain research, 166(3-4), 559–571. https://doi.org/10.1007/s00221-005-2396-5

Ayres, A. J., & Robbins, J. (2005). Sensory integration and the child: Understanding hidden sensory challenges: Western Psychological Services.

Barre, N., Morgan, A., Doyle, L. W., & Anderson, P. J. (2011). Language Abilities in Children Who Were Very Preterm and/or Very Low Birth Weight: A Meta-Analysis. The Journal of Pediatrics, 158(5), 766–774.e761. doi:https://doi.org/10.1016/j.jpeds.2010.10.032

Bergerbest, D., Ghahremani, D. G., & Gabrieli, J. D. (2004). Neural correlates of auditory repetition priming: reduced fMRI activation in the auditory cortex. Journal of cognitive neuroscience, 16(6), 966–977.

Bhutta, A. T., Cleves, M. A., Casey, P. H., Cradock, M. M., & Anand, K. (2002). Cognitive and behavioral outcomes of school-aged children who were born preterm: a meta-analysis. Jama, 288(6), 728–737.

Blackburn, S. (1998). Environmental impact of the NICU on developmental outcomes. Journal of pediatric nursing, 13(5), 279–289.

Brunet, D., Murray, M. M., & Michel, C. M. (2011). Spatiotemporal analysis of multichannel EEG: CARTOOL. Computational intelligence and neuroscience, 2011, 1–15.

Bucci, M. P., Wiener-Vacher, S., Trousson, C., Baud, O., & Biran, V. (2015). Subjective visual vertical and postural capability in children born prematurely. PloS one, 10(3), e0121616.

Carvalhais, C., da Silva, M., Xavier, A., Santos, J. (2017). Newborns safety at neonatal intensive care units: are they exposed to excessive noise during routine health care procedures?. Global Environment Health and Safety. 1. 1–3.

Chorna, O., Hamm, E., Shrivastava, H., & Maitre, N. L. (2018). Feasibility of event-related potential (ERP) biomarker use to study effects of mother’s voice exposure on speech sound differentiation of preterm infants. Developmental neuropsychology, 43(2), 123–134.

Chorna, O. D., Slaughter, J. C., Wang, L., Stark, A. R., & Maitre, N. L. (2014). A pacifier-activated music player with mother’s voice improves oral feeding in preterm infants. Pediatrics, 133(3), 462–468. https://doi.org/10.1542/peds.2013-2547

De Lucia, M., Tzovara, A., Bernasconi, F., Spierer, L., & Murray, M. M. (2012). Auditory perceptual decision-making based on semantic categorization of environmental sounds. Neuroimage, 60(3), 1704–1715.

De Lucia, M., Clarke, S., & Murray, M. M. (2010). A temporal hierarchy for conspecific vocalization discrimination in humans. Journal of Neuroscience, 30(33), 11210–11221.

Dimitrova, N., Turpin, H., Borghini, A., Harari, M. M., Urben, S., & Müller-Nix, C. (2018). Perinatal stress moderates the link between early and later emotional skills in very preterm-born children: an 11-year-long longitudinal study. Early human development, 121, 8–14.

Gallo, J., Dias, K. Z., Pereira, L. D., Azevedo, M. F. d., & Sousa, E. C. (2011). [Auditory processing evaluation in children born preterm]. Jornal da Sociedade Brasileira de Fonoaudiologia, 23, 95–101.

Gomot, M., Bruneau, N., Laurent, J.-P., Barthélémy, C., & Saliba, E. (2007). Left temporal impairment of auditory information processing in prematurely born 9-year-old children: an electrophysiological study. International journal of psychophysiology, 64(2), 123–129.

Guthrie, D., & Buchwald, J. S. (1991). Significance testing of difference potentials. Psychophysiology, 28(2), 240–244.

Husain, F. T., Horwitz, B. (2006). Experimental-neuromodeling framework for understanding auditory object processing: integrating data across multiple scales. J. Physiol. Paris 100, 133–141. doi: 10.1016/j.jphysparis.2006.09.006

Jackson, T. L., Ong, G. L., McIndoe, M. A., & Ripley, L. G. (2003). Monocular chromatic contrast threshold and achromatic contrast sensitivity in children born prematurely. American Journal of Ophthalmology, 136(4), 710–719.

Jansson-Verkasalo, E., Čeponien, R., Valkama, M., Vainionpää, L., Laitakari, K., Alku, P.,… Näätänen, R. (2003). Deficient speech-sound processing, as shown by the electrophysiologic brain mismatch negativity response, and naming ability in prematurely born children. Neuroscience letters, 348(1), 5–8.

Jansson-Verkasalo, E., Ruusuvirta, T., Huotilainen, M., Alku, P., Kushnerenko, E., Suominen, K.,… Tolonen, U. (2010). Atypical perceptual narrowing in prematurely born infants is associated with compromised language acquisition at 2 years of age. BMC neuroscience, 11(1), 88.

Johnson, S., & Marlow, N. (2011). Preterm birth and childhood psychiatric disorders. Pediatr Res, 69(5 Pt 2), 11R–18R.

Jongmans, M., Mercuri, E., Henderson, S., de Vries, L., Sonksen, P., & Dubowitz, L. (1996). Visual function of prematurely born children with and without perceptual-motor difficulties. Early human development, 45(1-2), 73–82.

Key, A. P., Lambert, E. W., Aschner, J. L., & Maitre, N. L. (2012). Influence of gestational age and postnatal age on speech sound processing in NICU infants. Psychophysiology, 49(5), 720–731.

Klaver, P., Latal, B., & Martin, E. (2015). Occipital cortical thickness in very low birth weight born adolescents predicts altered neural specialization of visual semantic category related neural networks. Neuropsychologia, 67, 41–54. https://doi.org/10.1016/j.neuropsychologia.2014.10.030

Knebel, J. F., & Murray, M. M. (2012). Towards a resolution of conflicting models of illusory contour processing in humans. Neuroimage, 59(3), 2808–2817.

Koenig, T., Kottlow, M., Stein, M., & Melie-García, L. (2011). Ragu: a free tool for the analysis of EEG and MEG event-related scalp field data using global randomization statistics. Computational Intelligence and Neuroscience, 2011, 4.

Koenig, T., Stein, M., Grieder, M., & Kottlow, M. (2014). A tutorial on data-driven methods for statistically assessing ERP topographies. Brain topography, 27(1), 72–83.

Korpilahti, P., Valkama, M., & Jansson-Verkasalo, E. (2016). Event-Related Potentials Reflect Deficits in Lexical Access: The N200 in Prematurely Born School-Aged Children. Folia Phoniatrica et Logopaedica, 68(4), 189–198.

Lehmann, D. (1987). Principles of spatial analysis. Handbook of electroencephalography and clinical neurophysiology: Methods of analysis of brain electrical and magnetic signals, 1, 309–354.

Lehmann, D., & Skrandies, W. (1980). Reference-free identification of components of checkerboard-evoked multichannel potential fields. Electroencephalography and clinical neurophysiology, 48(6), 609–621.

Lewis, J.W., Wightman, F.L., Brefczynski, J.A., Phinney, R.E., Binder, J.R., DeYoe, E.A.(2004).Human brain regions involved in recognising environmental sounds. Cereb Cortex, 14:1008–1021.

Lewis, J.W., Brefczynski, J.A., Phinney, R.E, Janik, J.J., DeYoe, E.A.(2005). Distinct cortical pathways for processing tool versus animal sounds. J Neurosci, 25:5148–5158.

Maitre, N. L., Key, A. P., Slaughter, J. C., Yoder, P. J., Neel, M. L., Richard, C., Wallace, M. T., & Murray, M. M. (2020). Neonatal Multisensory Processing in Preterm and Term Infants Predicts Sensory Reactivity and Internalizing Tendencies in Early Childhood. Brain topography, 33(5), 586–599.

Mahon, B. Z., Anzellotti, S., Schwarzbach, J., Zampini, M., & Caramazza, A. (2009). Category-specific organization in the human brain does not require visual experience. Neuron, 63(3), 397–405. https://doi.org/10.1016/j.neuron.2009.07.012

Maitre, N. L., Key, A. P., Chorna, O. D., Slaughter, J. C., Matusz, P. J., Wallace, M. T., & Murray, M. M. (2017). The dual nature of early-life experience on somatosensory processing in the human infant brain. Current biology, 27(7), 1048–1054.

Maitre, N. L., Slaughter, J. C., Aschner, J. L., & Key, A. P. (2014). Hemisphere differences in speech-sound event-related potentials in intensive care neonates: associations and predictive value for development in infancy. Journal of child neurology, 29(7), 903–911.

Mattioni, S., Rezk, M., Battal, C., Bottini, R., Cuculiza Mendoza, K.E., Oosterhof, N.N., Collignon, O. (2020) Categorical representation from sound and sight in the ventral occipito-temporal cortex of sighted and blind eLife 9:e50732 https://doi.org/10.7554/eLife.50732

Menendez, R. G. D. P., Andino, S. G., Lantz, G., Michel, C. M., & Landis, T. (2001). Noninvasive localization of electromagnetic epileptic activity. I. Method descriptions and simulations. Brain topography, 14, 131–137.

Menendez, R. G. D. P., Murray, M. M., Michel, C. M., Martuzzi, R., Andino, S. G. (2004). Electrical neuroimaging based on biophysical contstraints. Neuroimage, 21(2), 527–539.

Michel, C. M., & Murray, M. M. (2012). Towards the utilization of EEG as a brain imaging tool. Neuroimage, 61(2), 371–385.

Michel, C. M., Murray, M. M., Lantz, G., Gonzalez, S., Spinelli, L., & de Peralta, R. G. (2004). EEG source imaging. Clinical neurophysiology, 115(10), 2195–2222.

Mikkola, K., Wetzel, N., Leipälä, J., Serenius-Sirve, S., Schröger, E., Huotilainen, M., & Fellman, V. (2010). Behavioral and evoked potential measures of distraction in 5-year-old children born preterm. International journal of psychophysiology, 77(1), 8–12.

Murray, M. M., Michel, C. M., Grave de Peralta, R., Ortigue, S., Brunet, D., Gonzalez Andino, S., & Schnider, A. (2004). Rapid discrimination of visual and multisensory memories revealed by electrical neuroimaging. NeuroImage, 21(1), 125–135. https://doi.org/10.1016/j.neuroimage.2003.09.035

Murray, M. M., Foxe, J. J., & Wylie, G. R. (2005). The brain uses single-trial multisensory memories to discriminate without awareness. NeuroImage, 27(2), 473–478. https://doi.org/10.1016/j.neuroimage.2005.04.016

Murray, M. M., Brunet, D., & Michel, C. M. (2008). Topographic ERP analyses: a step-by-step tutorial review. Brain topography, 20(4), 249–264.

Murray, M. M., Camen, C., Andino, S. L. G., Bovet, P., & Clarke, S. (2006). Rapid brain discrimination of sounds of objects. Journal of Neuroscience, 26(4), 1293–1302.

Nevalainen, P., Pihko, E., Metsäranta, M., Andersson, S., Autti, T., & Lauronen, L. (2008). Does very premature birth affect the functioning of the somatosensory cortex?—A magnetoencephalography study. International journal of psychophysiology, 68(2), 85–93.

Paquette, N., Vannasing, P., Tremblay, J., Lefebvre, F., Roy, M.-S., McKerral, M.,… Gallagher, A. (2015). Early electrophysiological markers of atypical language processing in prematurely born infants. Neuropsychologia, 79, 21–32.

Perrin, F., Pernier, J., Bertnard, O., Giard, M., & Echallier, J. (1987). Mapping of scalp potentials by surface spline interpolation. Electroencephalography and clinical neurophysiology, 66(1), 75–81.

Pierrehumbert, B., Ramstein, T., Karmaniola, A. et al. Child care in the preschool years: Attachment, behaviour problems and cognitive development. Eur J Psychol Educ 11, 201–214 (1996). https://doi.org/10.1007/BF03172725

Pineda, R., Durant, P., Mathur, A., Inder, T., Wallendorf, M., Schlaggar, B.L. (2017). Auditory exposure in the neonatal intensive care unit: room type and other predictors. The Journal of Pediatrics, 183, 56–66.e3, https://doi.org/10.1016/j.jpeds.2016.12.072.

Pourtois, G., Delplanque, S., Michel, C., & Vuilleumier, P. (2008). Beyond conventional event-related brain potential (ERP): exploring the time-course of visual emotion processing using topographic and principal component analyses. Brain topography, 20(4), 265–277.

Retsa, C., Matusz, P. J., Schnupp, J. W., & Murray, M. M. (2018). What’s what in auditory cortices?. Neuroimage, 176, 29–40.

Retsa, C., Matusz, P. J., Schnupp, J. W. H., & Murray, M. M. (2020). Selective attention to sound features mediates cross-modal activation of visual cortices. Neuropsychologia, 144, 107498.

Salvari, V., Paraskevopoulos, E., Chalas, N., Muller, K., Wollbrink, A., Dobel, C., Korth, D., Pantev, C. (2019). Auditory categorization of manmade sounds versus natural sounds by means of MEG functional brain connectivity. Front. Neurosci., 13. DOI=10.3389/fnins.2019.01052

Saygın, A. P., Dick, F., W. Wilson, S. F., Dronkers, N., & Bates, E. (2003). Neural resources for processing language and environmental sounds: evidence from aphasia. Brain, 126(4), 928–945.

Spinelli, L., Andino, S. G., Lantz, G., Seeck, M., & Michel, C. M. (2000). Electromagnetic inverse solutions in anatomically constrained spherical head models. Brain topography, 13(2), 115.

Therien, J. M., Worwa, C. T., Mattia, F. R., & deRegnier, R. A. (2004). Altered pathways for auditory discrimination and recognition memory in preterm infants. Developmental medicine and child neurology, 46(12), 816–824. https://doi.org/10.1017/s0012162204001434

Tovar, D. A., Murray, M. M., & Wallace, M. T. (2020). Selective Enhancement of Object Representations through Multisensory Integration. The Journal of neuroscience: the official journal of the Society for Neuroscience, 40(29), 5604–5615. https://doi.org/10.1523/JNEUROSCI.2139-19.2020

Turpin, H., Urben, S., Ansermet, F., Borghini, A., Murray, M. M., & Müller-Nix, C. (2019). The interplay between prematurity, maternal stress and children’s intelligence quotient at age 11: A longitudinal study. Scientific reports, 9(1), 450.

Tzovara, A., Murray, M. M., Bourdaud, N., Chavarriaga, R., Millán, J. d. R., & De Lucia, M. (2012). The timing of exploratory decision-making revealed by single-trial topographic EEGanalyses. Neuroimage, 60(4), 1959–1969.

Vitale, F.M., Chirico, G., Lentini, C.(2021). Sensory Stimulation in the NICU Environment: Devices, Systems, and Procedures to Protect and Stimulate Premature Babies. Children, 8, 334. https://doi.org/10.3390/children8050334

Wachman, E. M., & Lahav, A. (2011). The effects of noise on preterm infants in the NICU. Archives of Disease in Childhood-Fetal and Neonatal Edition, 96(4), F305–F309.

Wickremasinghe, A., Rogers, E., Johnson, B., Shen, A., Barkovich, A., & Marco, E. (2013). Children born prematurely have atypical sensory profiles. Journal of Perinatology, 33(8), 631.

